# FREQ-NESS reveals age-related differences in frequency-resolved brain networks during auditory recognition and resting state

**DOI:** 10.1101/2025.06.30.662094

**Authors:** Chiara Malvaso, Gemma Fernández-Rubio, Mattia Rosso, Elisa Serra, Vera Rudi, Peter Vuust, Morten L. Kringelbach, Claudia Testa, Leonardo Bonetti

## Abstract

Understanding how brain networks operate across different frequencies during cognitive tasks, and how these dynamics change with age, remains a central challenge in cognitive neuroscience. While previous studies have focused on resting-state activity and passive listening, less is known about frequency-specific brain dynamics during event-related tasks that require active memory engagement. In this study, we extend the recently developed FREQ-NESS analytical pipeline by adapting it to event-related task and resting state source-reconstructed magnetoencephalography (MEG) data from 70 healthy participants. This method quantified the variance explained by frequency-specific brain networks, their spatial organization, and associated time-resolved power estimates. We found significant effects of age, condition, and their interaction in the variance explained by leading components at 1.07 Hz, 2.86 Hz, and 10.00 Hz. Younger adults exhibited stronger peaks at 1.07 and 2.86 Hz during the task and a more pronounced 10.00 Hz peak at rest, whereas older adults showed the opposite pattern. Time-frequency analysis revealed age- and condition-dependent desynchronization in the alpha and beta bands (7.10–22.90 Hz). These findings demonstrate the effectiveness of the adapted FREQ-NESS pipeline for event-related tasks and highlight the importance of frequency-resolved network analysis for characterizing age-related changes in active auditory memory processing.

## 1 Introduction

The natural aging process is characterized by a progressive decline in memory, attention, and other cognitive functions [1, 2, 3, 4], while the ability to perform activities of daily living generally remains preserved [5]. As global life expectancy continues to increase, driven by advancements in healthcare and living conditions, the investigation of both healthy and pathological aging has gained critical importance. Neurophysiological measures have become particularly valuable in this regard, offering insights into the neural mechanisms underlying age-related cognitive changes [6]. A widely used experimental condition in such studies is the resting awake state with eyes closed, as it provides a standardized, cost-effective protocol that is well suited for neurophysiological recordings in older adults [7]. Numerous resting-state studies have reported that both pathological and healthy aging were associated with characteristic spectral “slowing” in the power spectrum [8, 9, 10, 11, 12]. This was particularly evident in case of pathological aging where patients were diagnosed with mild cognitive impairment (MCI) and dementia [8, 9]. This slowing typically manifests as a reduction in both absolute and relative power within the higher frequency bands (alpha and beta, 8–30 Hz), accompanied by an increase in power within the lower frequency bands (delta and theta, 0.5–8 Hz) [13, 14]. Additionally, reductions in the individual alpha frequency (IAF) peak have been observed in patients with Alzheimer’s disease dementia (ADD) and MCI, relative to healthy controls [15, 16]. A similar pattern was also found in healthy aging, with older adults displaying lower alpha power and a reduced IAF peak compared to younger individuals [10, 11, 12].

Resting-state recordings with eyes closed during quiet vigilance have become the standard approach in EEG and MEG studies of both healthy and pathological aging, largely due to their ease of implementation [6]. However, in comparison to the eyes-open condition, the eyes-closed state is associated with increased alpha power [17], which may influence the interpretation of findings related to power levels and resting-state brain dynamics [18]. Notably, even though they are less common, studies using eyes-open restingstate paradigms have reported similar age-related spectral changes, specifically a general “slowing” of the power spectrum. These findings suggest that the observed age-related alterations in oscillatory dynamics are not solely specific to the eyes-closed condition. Nonetheless, resting-state paradigms capture only background neural activity and do not engage brain processes specific to cognitive functions such as memory or attention, which are critical to understanding both healthy and pathological aging. To complement and extend the findings derived from resting-state analyses, task-based approaches are also employed in aging research. Among the various available paradigms, auditory and visual oddball paradigms are among the most frequently used [6]. Studies utilizing the oddball paradigm have reported attenuated mismatch negativity responses, indicative of automatic memory processing, in individuals with ADD, vascular dementia (VD), and MCI, when compared to control groups [19, 20, 21], as well as in healthy older adults relative to younger individuals [22, 23]. In addition to investigating automatic auditory and visual change detection through oddball paradigms, tasks evaluating working memory, inhibitory control, and short-term memory are also among the most commonly employed most commonly employed, contributing to the understanding of the neural mechanisms underlying cognitive decline in aging populations [6].

Using these paradigms, previous studies showed age-related changes in event-related potentials/fields (ERP/F). For example, several studies have demonstrated that older adults, compared to younger adults, exhibit a significantly greater reduction in N400 amplitude, as well as delayed peak and/or onset latency during semantic processing and sentence comprehension tasks [24, 25, 26, 27]. Furthermore, across various tasks—including attention paradigms, response inhibition tasks, and spatial processing assessments—reduced P300 amplitudes have been consistently observed in healthy older adults compared to their younger counterparts [22, 28, 29]. Complementary to ERP/F analyses, investigations of event-related oscillations (EROs) and event-related desynchronization (ERD) have exhibited substantial variability across studies, likely reflecting differences in task design. Nonetheless, age-related increases in movement-related beta desynchronization and beta bursts during motor and sensory processing tasks have been reliably reported [30, 31, 32]. A recent study showed a dissociation between ERPs and additional properties of the signal. Criscuolo and colleagues [33] showed that older adults presented larger ERPs but educed phase-coherence at the stimulation frequency in a simple task where participants were presented with isochronous auditory sequences.

While providing valuable insights, the aforesaid tasks predominantly rely on static stimuli, such as images or numerical data. To advance our understanding of age-related changes, a promising direction involves tasks requiring time-dependent information processing, thereby enabling the investigation of how information is handled over time. Among emerging approaches, music-based tasks have recently gained attention as a promising means to investigate predictive brain mechanisms within the context of long-term memory encoding and recognition [34, 35, 36, 37, 38, 39, 40, 41, 42, 43, 44, 45]. In the context of age-related studies, one investigation reported that older adults exhibit a reorganization of functional brain activity during the recognition of previously memorized sequences, in comparison to younger adults [46]. Specifically, the study found increased early activity in sensory regions, such as the left auditory cortex, alongside only moderate reductions in activity within the medial temporal lobe and prefrontal regions. In response to varied sequences, older adults demonstrated a pronounced reduction in fast-scale functionality across higher-order brain regions, including the hippocampus, ventromedial prefrontal cortex, and inferior temporal cortices, whereas no differences were observed in the auditory cortex [46]. The aforementioned study focused on broadband signal analysis. To complement and extend those findings, the present study performs a narrowband analysis on the same dataset, with the aim of identifying age-related differences within specific frequency bands. In particular, this work expands on the FREQ-NESS analytical pipeline, presented in [47],which was designed to estimate both the activation and spatial configuration of concurrent brain networks across frequencies. This approach analyzes the frequency-resolved multivariate covariance of whole-brain voxel time series using Generalized Eigenvector Decomposition (GED) [48], a linear decomposition technique applied to source-reconstructed MEG data acquired during resting state and isochronous auditory stimulation. GED enhances the contrast between narrowband and broadband signals by acting as a spatial filter that isolates task-relevant neural patterns while enabling dimensionality reduction [49, 50, 51, 52, 53]. Owing to their mathematical tractability and capacity to integrate signals across multiple channels via weighted summation, linear methods are particularly suitable for for multivariate analysis and brain networks estimation decoding studies [54, 47]. In particular, FREQ-NESS relies on the application of GED on source-reconstructed MEG data, presenting several advantages: it imposes no spatial or anatomical constraints and is invariant to the order and spatial configuration of data channels (e.g., electrodes, sensors, pixels, or voxels), enabling physiologically meaningful interpretations [54] without reliance on predefined anatomical models. Additionally, this method is deterministic and non-iterative, ensuring reproducible results across repeated analyses. Its computational efficiency is notable, with most processing time attributed to preprocessing steps such as temporal filtering.

The present study builds on the methodology introduced in [47] by applying the FREQ-NESS analysis pipeline to MEG data recorded during a melody recognition task, , aiming to identify frequency-specific differences across brain networks. FREQ-NESS [47] provides information on the relative significance of each identified network, as the variance explained by each component reflects its importance, projections along the identified directions allow for the investigation of temporal dynamics, and the associated weights yield activation patterns that localize brain regions involved in frequency-specific processing. In doing so, the study seeks to address the previously identified gap in aging research by complementing the analysis conducted in [46], specifically by investigating differences between younger and older adults frequency specific brain networks

## 2 Methods

### 2.1 Participants

The dataset used in the present study is the same as that employed in [46], originally comprising 76 participants after the exclusion of one individual who did not complete the experimental task. However, in the current analysis, an additional six participants whose task accuracy fell below 50% were excluded, resulting in a final sample of 70 participants. Based on self-reported biological sex (distinct from gender identity, which was not assessed as it was not pertinent to the study objectives), the sample included 31 males and 39 females. Participants were arranged into two age-based sets: younger adults (*n* = 37; 18 females, 19 males) and older adults (*n* = 33; 21 females, 12 males). The younger group ranged in age from 18 to 25 years (21.89 ± 2.05), and the older one ranged from 60 to 81 years (66.61 ± 5.02).

**Figure 1:**
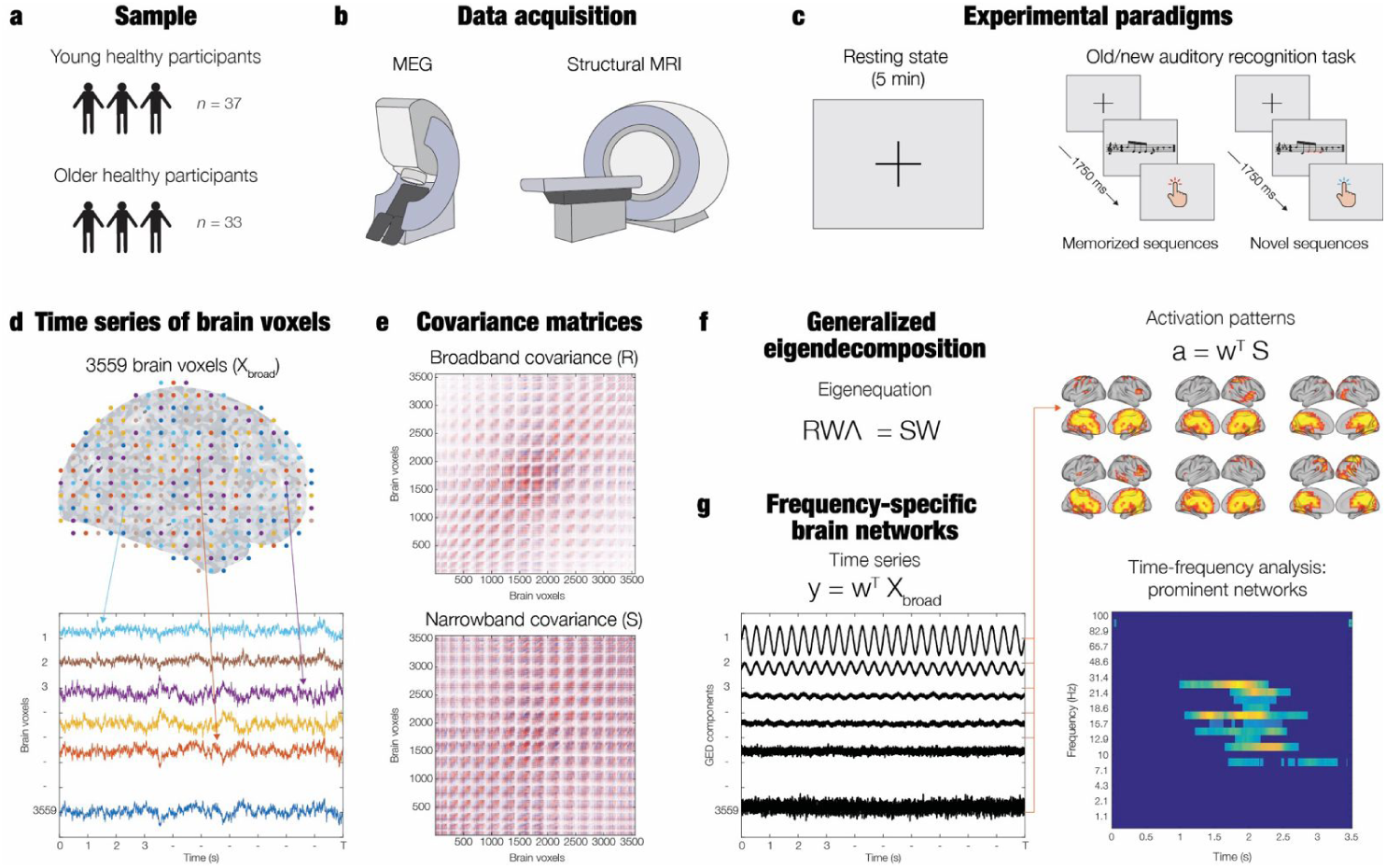
Overview of the methodology employed in the present study.a – The final sample consisted of 70 healthy participants divided into two groups: younger adults (*n* = 37) and older adults (*n* = 33). b – MEG data were recorded to obtain the participants’ brain activity. MRI data were collected for co-registration purposes. c – MEG recordings were obtained during both resting state and task performance. During the melody recognition task, participants classified five-tone auditory sequences, presented in random order, as either “old” (memorised sequences, M) or “new” (novel sequences, N) via button presses. d – Source reconstruction was performed using a beamforming algorithm, resulting in time series data for 3559 brain voxels based on an 8-mm grid brain parcellation. e – Covariance matrices were computed from the voxel time series data for both broad-band (**R**) and narrow-band (**S**) signals. f – Generalized Eigenvector Decomposition (GED) was performed by solving the equation **RWΛ** = **SW**; the weight matrix **W** maximizes the contrast between **R** and **S**, and the eigenvalues **Λ** quantify the variance explained by each network component. g – The weights **W** were used to project the voxel time series into the GED-defined component space via **y** = **w***^T^* **X***_broad_*, and to compute the corresponding spatial activation patterns using **a** = **w***^T^* **S**. A Morlet wavelet transform was then applied to the resulting time series **y** to extract power time series for time-frequency analysis.

All participants were of Danish nationality. Ethical approval for the study was granted by the Institutional Review Board at Aarhus University (Case No. DNC-IRB-2021-012), and all procedures conformed to the ethical standards of the Declaration of Helsinki. Informed consent was obtained from each participant prior to participation. Participants were compensated for their involvement, and all procedures were carried out in accordance with applicable ethical guidelines governing research involving human subjects.

### 2.2 Experimental design

Participants were presented with a melody recognition paradigm based on an old/new recognition task, which has been extensively employed in previous studies ([55, 56, 57, 58, 59]). Simultaneously, participants’ brain activity was recorded using magnetoencephalography (MEG). They listened twice to a short musical excerpt, approximately 25 seconds long, and were instructed to memorize it as accurately as possible. This excerpt consisted of the first four measures of Johann Sebastian Bach’s Prelude No. 2 in C Minor, BWV 847. The audio file used in the experiment was a wave format generated with Finale (version 26, MakeMusic, Boulder, CO) and played through Psychopy v3.0. The volume of the musical stimulus was set at 60 dB for most participants (n = 67), while for nine participants over the age of 70—who exhibited very mild, age-related hearing impairments—the volume was adjusted to an average of 70 dB. To minimize the need for individual volume adjustments, the audio was designed to predominantly include frequencies between 125 and 650 Hz, a range that is only slightly affected by typical age-related hearing loss. Each tone in the piece had a uniform duration of about 350 milliseconds. During the second phase of the task, participants were presented with 81 five-tone musical sequences, each lasting 1750 milliseconds. For each sequence, they were asked to indicate whether it was taken from the original piece (memorized or “old” sequence [M]) or if it was a different sequence altogether (novel or “new” sequence [N]). As stated above, participants who could not identify correctly at least 50% of the presented sequences were excluded.

### 2.3 Data acquisition

MEG recordings were acquired at Aarhus University Hospital (AUH), Aarhus, Denmark, using a 306-channel Elekta Neuromag TRIUX MEG system. The data were acquired with analog filtering in the range of 0.1–330 Hz, at a sampling rate of 1000 Hz. For accurate coregistration with anatomical MRI scans, participants’ head shapes and the locations of four Head Position Indicator (HPI) coils were digitised using a 3D Polhemus Fastrak system (Colchester, VT, USA). To facilitate later removal of physiological artifacts, two sets of bipolar electrodes were used during MEG acquisition to monitor cardiac activity (ECG) and eye movements (EOG).

Structural MRI scans were obtained on a CE-approved 3 T Siemens scanner at AUH, using a T1-weighted sequence with the following parameters: echo time (TE) = 2.61 ms, repetition time (TR) = 2300 ms, matrix size = 256 × 256, echo spacing = 7.6 ms, and bandwidth = 290 Hz/Px, resulting in a 1×1×1 mm spatial resolution. MEG and MRI recordings were performed on separate days.

### 2.4 Pre-processing

MEG recordings were acquired from 204 planar gradiometers and 102 magnetometers. Initial pre-processing was conducted using MaxFilter [60] (version 2.2.15) to attenuate external interferences.The following parameters were used within MaxFilter: spatiotemporal signal space separation (SSS), downsampling from 1000 Hz to 250 Hz, a correlation limit of 0.98 between inner and outer subspaces to reject overlapping signals during spatiotemporal SSS, and movement compensation based on continuous head position indicator (cHPI) coils with a default step size of 10 ms. The data were converted into Statistical Parametric Mapping (SPM) format and subsequently analyzed in MATLAB, employing both custom-developed code (LBDP) and the freely available Oxford Centre for Human Brain Activity (OHBA) Software Library (OSL) [61], which integrates the Fieldtrip[62], FSL (version 6.0)[63], and SPM12[64] toolboxes. Visual inspection of the filtered MEG data using OSLview confirmed the successful removal of prominent artifacts, which constituted less than 0.1% of the total dataset. Eyeblink and heartbeat interference were eliminated using Independent Component Analysis (ICA) [65]. This process involved decomposing the original signal into independent components, removing those associated with eyeblink and heartbeat activity, and reconstructing the signal from the remaining components. The resulting signal was then segmented into 81 trials, followed by baseline correction performed by subtracting the mean baseline signal from the post-stimulus brain response. Each trial had a duration of 3500 ms (comprising 3400 ms post-onset of the first tone in the musical sequence and 100 ms of baseline). The trials were evenly distributed into three categories (M, NT1, NT3), with 27 trials per group.

### 2.5 Source reconstruction

MEG recordings offer insights into neural activity outside the head but do not directly reveal the specific brain sources underlying this activity. To estimate the brain sources responsible for the recorded signals, we employed a source reconstruction protocol that integrated custom-developed code along with the OSL, SPM, and FieldTrip toolboxes. The reconstruction process consisted of two main steps: (*i*) designing a forward model, and (*ii*) computing the inverse solution. The forward model was constructed in the first step, utilizing a single-shell model with an 8-mm grid. In this model, each brain source was represented as an active dipole, with the model describing how the activity of these dipoles would be detected by the MEG sensors. Magnetometer channels and an 8-mm grid were used to generate 3559 dipole locations throughout the entire brain, corresponding to individual voxels. To ensure accurate alignment of the MEG data with the brain’s anatomy, the data were co-registered with individual T1-weighted MRI scans using 3D digitizer information, which included landmarks such as the nose and the left and right pre-auricular points. The forward model was computed using the single-shell method, resulting in a leadfield model represented by the matrix **L** with dimension (sources × MEG channels)[66]. In instances where individual structural T1-weighted images were unavailable, the MNI152-T1 template with 8-mm spatial resolution was used for leadfield computation. The leadfield model was calculated for the three principal orientations of each brain source (dipole), as is standard practice[66]. To simplify the beamforming output, these orientations were subsequently reduced to a single effective orientation using the singular value decomposition (SVD) algorithm, applied to the matrix product:

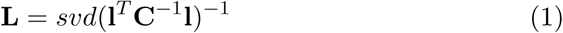

Here, **l** denotes the leadfield model encompassing the three orientations, while **L** represents the resulting single-orientation model used for source reconstruction. This procedure is commonly employed to simplify the beamforming output[67]. In the second step, a beamforming algorithm was utilized as the inverse model. This approach assigns weights to individual source locations (dipoles) in order to isolate the contribution of each brain source to the recorded MEG signal. The algorithm was applied at each time point of the recording, thereby enabling the reconstruction of the spatial distribution of MEG activity over time. Specifically, for every time point, the beamformer computes a distinct set of weights to project the MEG sensor data onto the corresponding active dipoles. The procedure for computing the inverse solution is described as follows. The MEG signal recorded at a given time point, denoted by **B**, is mathematically related to the underlying neuronal activity through the following expression:

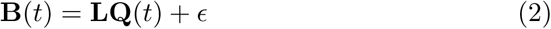

Here, **Q**(*t*) denotes the strength of neuronal activity, **B**(*t*) is a column vector containing the magnetic field measurements at time *t*, **L** is the lead field, and *ɛ* represents the noise[68]. Solving the inverse problem requires computing **Q**(*t*). Within the beamforming framework, this involves calculating a set of weights that are applied to the MEG sensor data at each time point. This operation is repeated for each individual dipole **q**(*t*), as described by:

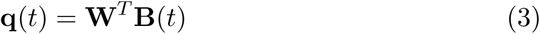

where the superscript *T* indicates matrix transposition, and **W** represents the set of weights to be computed. The beamforming method relies on the product of the leadfield matrix **L** and the covariance matrix **C**, which captures the covariance between MEG sensors. This covariance matrix is estimated from the signal obtained by concatenating all experimental trials. Specifically, for each brain source (dipole) **q**, the weights **W***_q_* are computed as:

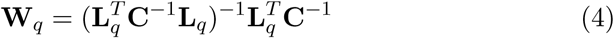

The computed weights were then applied across all brain sources and time points. The covariance matrix **C** was derived from the continuous signal resulting from the concatenation of all experimental trials. To mitigate reconstruction bias toward the center of the head, the weights were normalized according to the approach described. Given that the analysis focused on evoked responses, the weights were applied to neural activity averaged over trials. This procedure yielded a time series for each of the 3559 brain sources across all experimental conditions. To resolve the sign ambiguity inherent to evoked response time series, the polarity of each source was matched to the N100 response elicited by the first tone in the auditory sequences [55, 56, 57, 58, 59]. The source-reconstructed time series were provided as input to the FREQ-NESS analytical pipeline, originally presented in [47] and adapted in the current work for event-related tasks.

### 2.6 Filtering

Given that the primary aim was to investigate potential differences across various frequencies - while leaving a comprehensive, systematic exploration of all possible frequencies to future work - 28 frequencies of interest were selected within the range of 0 to 100 Hz. Due to the nature of the stimulus, the stimulation frequency remained approximately constant at 2.857 Hz. Accordingly, the frequency set included both harmonics and subharmonics of the stimulation frequency. The selection strategy was guided by the expectation that the most prominent differences would emerge below approximately 20 Hz. Therefore, a denser sampling of frequencies was applied within this lower range, with progressively sparser spacing at higher frequencies.

To isolate frequency-specific components, a Gaussian filter was utilized. This filter functions by convolving the input signal with the impulse response of a Gaussian function [69]. Since the filter operates in the frequency domain, it was necessary to specify both the central frequency and the standard deviation of the Gaussian envelope in that domain. The filter amplitude varied with frequency: a narrower filter (i.e., smaller standard deviation) was used at lower frequencies, while the standard deviation increased linearly with the central frequency for frequencies above 2.86 Hz. For frequencies below the stimulation frequency, the amplitude was maintained constant to avoid distortion of the frequency response caused by excessive attenuation.

### 2.7 Resting state epoching

As outlined in the pre-processing section (2.4), MEG listening data consisted of a series of trials, each lasting 3500 ms. Of this duration, the first 100 ms served as a baseline, followed by 3400 ms post-onset of the initial tone in the musical sequence. In the present analysis, FREQ-NESS was applied independently to each trial. While data from the listening condition were epoched during preprocessing—thereby isolating individual trials—epoching the resting-state data posed a challenge due to the absence of event-related triggers, such as stimulus onsets, that could define epoch boundaries.

To address this, an artificial epoching procedure was developed based on the following criteria: (*i*) each epoch was fixed at a duration of 3500 ms, matching the listening condition; (*ii*) epochs were non-overlapping; and (*iii*) the total number of epochs in the resting-state data equaled the number of trials in the corresponding listening condition for each subject. This was achieved by dividing the continuous resting-state recording into a number of equal intervals corresponding to the number of trials. Within each interval, a random starting time point was selected, ensuring that the subsequent 3500 ms did not overlap with the next interval.

This preliminary segmentation ensured that epochs were uniformly distributed across the entire resting-state recording. By constraining the random selection within predefined intervals, this method mitigated the risk of temporal clustering—i.e., the concentration of epochs in specific portions of the recording—thus promoting a more representative sampling of the resting-state data.

### 2.8 Generalized Eigenvector Decomposition

The goal of Generalized Eigendecomposition (GED) is to identify a set of channel weights that maximize the contrast between a signal and a reference by enhancing discriminative features while suppressing shared covariance patterns. Specifically, inter-channel covariance structures common to both the signal (**S**) and the reference (**R**) are disregarded, allowing the method to emphasize components that distinguish the signal from the background.

The channel weight vector corresponding to the largest eigenvalue is used as a spatial filter. Applying this filter to the multichannel time series yields a component time series that optimally enhances contrast according to the covariance structure of the data used to compute **S** and **R**. In the present analysis, GED was applied to source-reconstructed brain activity to isolate spatial networks operating at specific frequencies. For each frequency of interest, the method identified the weighted combination of voxels that best differentiated narrow-band oscillatory activity from broadband background activity by maximizing the contrast between their covariance matrices.

GED computes the set of eigenvectors **W** and eigenvalues **Λ** that contain the voxel weights maximizing the separation between the signal covariance (**S**) and the reference covariance (**R**). These covariance matrices were estimated in a single step through matrix multiplication of mean-centered voxel data, separately for the narrow-band and broadband signals, as follows:

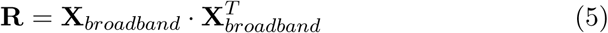

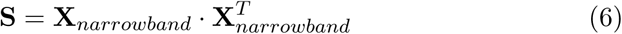

The computation described in equation 6 was repeated for each selected frequency. Since maintaining the frequency-resolved content of the signal requires avoiding averaging operations, which would suppress relevant spectral information [70], covariance matrices were computed separately for each subject and each trial.

The matrix of weights **W** was obtained by solving the following equation:

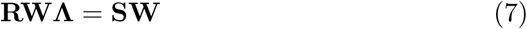

Here, **Λ** denotes the eigenvalues matrix, which quantifies the relative importance of each component in terms of the variance it explains. The corresponding eigenvectors were used as spatial filters; each was transposed and then multiplied by the broadband voxel data matrix to reconstruct the time series of the target signal components:

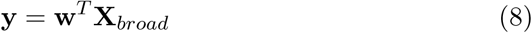

Subsequently, the spatial projection of the components in voxel space was computed. This was achieved by multiplying each individual raw weight vector **w** with the covariance matrix **S**. The absolute value of the resulting product was then taken to determine the absolute voxel activation strengths associated with the estimated brain network [47]. The resulting topographical distribution of the components was interpreted as the network activation patterns **a**:

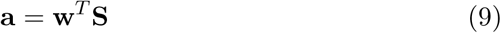

To improve the numerical stability of GED in cases of rank deficiency, regularization was applied to both covariance matrices by adding a small constant to their diagonals. Regularization consists of introducing a constant into the cost function of an optimization algorithm [48]. The regularization value was pre-determined and applied uniformly across all covariance matrices [48]. Specifically, a constant value of 10*^−^*^6^ was added to **S**, while for **R**, the regularization term corresponded to 1% of its average eigenvalue [47].

### 2.9 Frequency Analysis

To investigate potential frequency-resolved differences associated with the experimental conditions (auditory memory task or resting state) and the participants’ age (young and older groups), an analysis in the frequency domain was performed. This was accomplished by examining the participants’ group and experimental condition in relation to both the variance explained by each component and the induced responses, which were computed for each component using a Morlet Wavelet Transform.

#### 2.9.1 Variance explained Analysis

The initial phase of the analysis aimed to identify significant differences in the variance explained by the first component across frequencies, considering both experimental conditions and age groups. The first component was selected based on prior justification: as previously discussed, the eigenvector derived from equation 7 reflects the relative importance of each component. Consequently, the first component accounts for the greatest proportion of variance and thus captures the most relevant dynamics in the dataset. To assess the presence of significant effects attributable to age, condition, or their interaction, an Analysis of Variance (ANOVA) was performed. The analysis was conducted using the MATLAB function *anovan*, which performs a

multiway (n-way) ANOVA.

#### 2.9.2 Morlet Wavelet Transform

The second analytical stage involved applying the Morlet Wavelet Transform to the time series data of each subject [71].

Following the methodology outlined in [72], the wavelet was constructed by applying a Gaussian envelope with a defined full width at half maximum (FWHM), followed by the Hilbert Transform. This procedure was independently applied to each subject’s time series and repeated for both the task and resting conditions.

The analysis proceeded as follows:

1. For each frequency of interest, a sequence of operations was performed to generate a matrix representing the induced neural responses:

(i) A Gaussian filter, parameterized by the desired amplitude and FWHM, was applied to the continuous time series, independent of trial segmentation.

(ii) The filtered signal was then subjected to the Hilbert Transform, and the squared modulus of the resulting complex signal was computed.

(iii) Power was normalized using the formula:

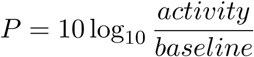

where *activity* denotes the post-stimulus induced response of the GED components, and *baseline* refers to the pre-stimulus interval.

(iv) The power time series was averaged across trials to obtain a representative induced response for each frequency.

2. The frequency-specific induced responses were concatenated to form a two-dimensional matrix with dimensions (frequency, time).

3. This matrix was then normalized by dividing each element by the matrix’s maximum value.

Subsequently, an ANOVA test was applied to the induced responses of the GED components. This test was conducted independently at each time point and frequency. The input to the ANOVA consisted of a matrix containing the induced responses for each subject under both experimental conditions (auditory memory task and resting). The goal was to identify statistically significant effects attributable to age, condition, and their interaction. To address the issue of multiple comparisons, a cluster-based permutation test was employed. Clusters were defined within a binary 2D matrix, and their statistical significance was evaluated using Monte Carlo simulations with 100 permutations.

## 3 Results

### 3.1 Overview on experimental design and analysis

In the current study, the FREQ-NESS pipeline described in [47] was utilized, with modifications tailored to the context of event-related analysis. This pipeline employs Generalized Eigenvector Decomposition (GED) to examine frequency-specific differences related to age, as well as distinctions between task and resting-state conditions. Magnetoencephalography (MEG) recordings were collected from 70 participants, divided into two age groups (younger and older adults), under two experimental conditions: resting state and a melody recognition task using an old/new paradigm [55, 56, 57, 58, 59].

FREQ-NESS was applied to characterize both temporal and spatial differences in brain networks operating at specific frequencies across age groups and experimental conditions. This was achieved by performing GED on the full brain voxel data matrix, reconstructed via beamforming. The procedure was repeated for 28 frequencies of interest, spanning the range from 0 to 100 Hz.

GED yields a set of eigenvectors that decompose the multivariate signal into components representing the most prominent narrowband activity at each frequency, contrasted against broadband activity. The associated eigenvalues quantify the strength of this contrast, indicating how much more variance is present in the narrowband signal relative to the broadband reference along each eigenvector-defined direction. The resulting weights were used to project the data along the direction of maximum variance—defined by the eigenvector associated with the highest eigenvalue—thus extracting the time series of the target component. These same eigenvectors were also used to filter the narrowband covariance matrix, enabling computation of the spatial activation patterns corresponding to each component.

### 3.2 Frequency-specific brain networks and age

The initial stage of the analysis focused on the variance explained by the first component, with the objective of identifying significant differences across frequencies as a function of experimental condition and age group. To evaluate these effects, an ANOVA test was employed. This analysis revealed three frequencies at which statistically significant differences were observed: 1.07 Hz, 2.86 Hz, and 10.00 Hz. F-values and p-values obtained from the ANOVA test that resulted significant for at least one source of variation (condition, age or their interaction) are schematically reported in table 1. Figure 2 displays the variance explained by the first components, as well as the brain activation patterns associated with the frequencies identified as significant.

**Figure 2:**
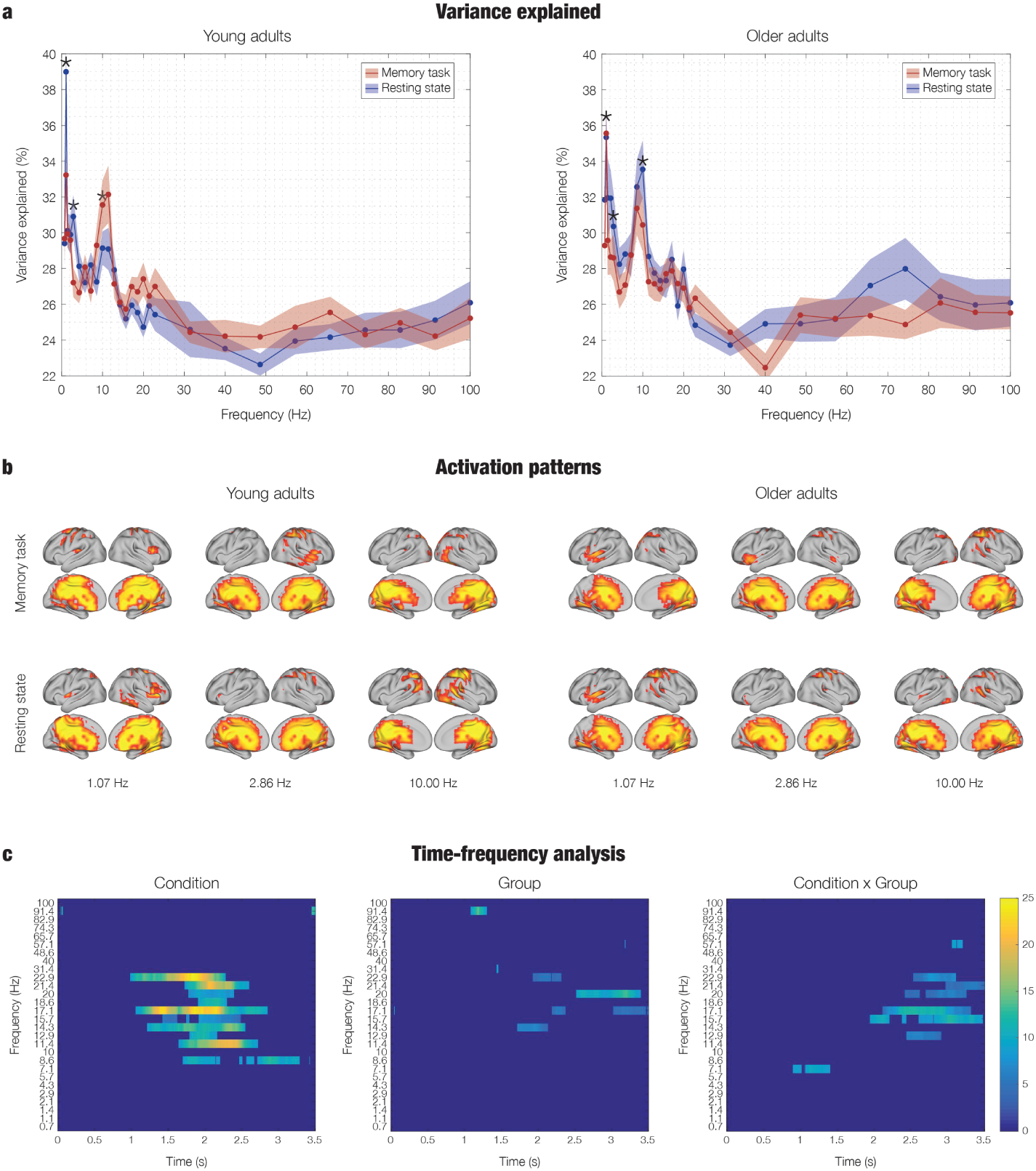
Summary of the results. a – Variance explained by the first component obtained through FREQ-NESS, shown for young adults (left panel) and older adults (right panel). For both groups, data are presented for resting state (red line) and task (blue line). Asterisks (∗) indicate frequencies at. The plotted lines indicate the mean variance explained across subjects, while the shaded regions denote the standard error of the mean. b – Spatial activation patterns for young (left) and older (right) adults corresponding to the networks at the three frequencies identified as significant. c – Results of the statistical analysis on the power time series. The plot displays F-values corresponding to significant effects of condition, age, or their interaction. F-values associated with non-significant effects were set to 0 to enhance the clarity of the visualization.

**Table 1:** Results of the ANOVA test conducted on the first component of the variance explained, derived from the GED computation with covariance on single trials. Only frequencies exhibiting a significant effect in at least one source of variation are reported.

| Frequency [Hz] | Source of Variation | F-value | p-value |
| --- | --- | --- | --- |
| 1.07 | *Age* | 0.54 | 0.46 |
|  | *Condition* | 4.05 | 0.05 |
|  | *Age\*Conditon* | 4.22 | 0.04 |
| 2.86 | *Age* | 0.19 | 0.66 |
|  | *Condition* | 6.63 | 0.01 |
|  | *Age\*Conditon* | 0.22 | 0.64 |
| 10.00 | *Age* | 1.47 | 0.23 |
|  | *Condition* | 0.23 | 0.80 |
|  | *Age\*Conditon* | 4.15 | 0.04 |

At 1.07 Hz, the ANOVA indicated significant effects for both the Condition factor (*p* = 0.05) and the interaction between Age and Condition (*p* = 0.04). Correspondingly, the variance explained by the first component exhibited a more prominent peak at this frequency in the task condition compared to the resting state among younger participants. In contrast, for the older group, the effect was less distinct, with peaks present in both conditions.

The second frequency identified as significant, 2.86 Hz, corresponds directly to the stimulation frequency. The ANOVA test revealed a significant effect of condition at this frequency, with *p*= 0.01.

The third frequency identified as significant by the ANOVA test was 10.00 Hz, with the effect attributed to the interaction between age and condition (*p*= 0.04). Among younger adults, this frequency corresponds to a distinct peak in the variance explained during the resting state. In contrast, older adults exhibited a similar peak at slightly lower frequencies during rest.

### 3.3 Brain networks time-frequency analysis

The second analytical procedure involved computing the Morlet wavelet transform of the time series for each subject. This transformation yielded a matrix of dimensions (frequency, time) per subject, capturing the temporal evolution of spectral power across the frequency spectrum. Each element of this matrix was subsequently subjected to an ANOVA test to assess the presence of statistically significant effects at each time point and frequency, with respect to age, condition, or their interaction.

The results are presented in Figure 2. To enhance the clarity of the resulting visualizations, F-values corresponding to non-significant effects—determined via a permutation test with correction for multiple comparisons—were set to zero. This masking procedure allowed for a more focused depiction of the specific time-frequency regions exhibiting statistically significant differences. The scale of the color bar axis is identical across all three plots to facilitate a more effective visual inspection. It should be noted that, the wavelet transform was computed independently for each of the 28 frequencies of interest, and the results were subsequently concatenated. Consequently, the values on the y-axis of the plots in this section correspond to the aforementioned frequencies. Since these frequencies are not equally spaced, but rather concentrated at lower values (below 20 Hz), a linear scale ranging from 0 to 100 on the y-axis would not accurately represent the actual frequency values used.

The analysis of condition-related effects revealed a significant difference between the resting and auditory task states within the frequency range of 7.10 to 22.90 Hz, encompassing the alpha and beta bands, following stimulus onset.

The analysis of the age factor revealed fewer significant time-frequency points and lower F-values compared to the condition-related analysis. In contrast, the interaction between age and condition produced more substantial results. Both analysis revealed significant effects within the 12.90–22.90 Hz range, with additional significant differences appearing at lower frequencies within the alpha band.

## 4 Discussion

In the present study, the FREQ-NESS method introduced in [47] was utilized to investigate temporal and spatial differences in brain networks operating at specific frequencies across age groups (young versus older adults) and experimental conditions (auditory memory task versus resting state with open eyes). In contrast to the study reported in [47], which concentrated on resting state and passive listening, the current work applied the same approach to investigate brain networks elicited during an event-related task. As results from the FREQ-NESS pipeline introduced by [47], in the current study the variance explained by the GED components at each frequency exhibited a steep exponential decay. When eigenvalues are ordered from highest to lowest, this pattern indicates that the first component accounts for the majority of the explained variance. Consequently, the current study concentrated exclusively on the first component, based on the premise that the remaining components reflect less relevant data dynamics. The variance explained by the first network at each frequency was significantly different for young and older participants as well as for resting and task conditions, at three frequencies: 1.07, 2.86, and 10.00 Hz, as determined by an ANOVA test. At 1.07 Hz, younger adults exhibited a peak in the variance explained during the listening condition, in contrast to older partcipants, for whom the explained variance at this frequency remained comparable across both task and resting states. A similar pattern was observed at 2.86 Hz, where younger participants showed a pronounced peak during task, which was less evident in older individuals. At 10.00 Hz, younger adults demonstrated a peak in the variance explained during the resting state, whereas in older adults, the corresponding peak during rest occurred at slightly lower frequencies.

Time-frequency analysis of the most prominent frequency-resolved brain networks indicated a significant distinction between resting and task conditions in frequency bands ranging from 7.10 to 22.90 Hz, encompassing the alpha and beta ranges, following stimulus onset. The main effect of age yielded fewer significant data points and lower F-values relative to the previous analysis. In contrast, the interaction between age and condition produced more prominent effects. In both cases, significant results were observed within the 12.90–22.90 Hz range, with additional significant differences appearing at lower frequencies within the alpha band.

As anticipated, the results of the present study are consistent with those reported in [46], which utilized the same dataset, albeit with minor differences in the final samples used for the analysis. That prior work identified a substantial reorganization of brain function associated with aging in the context of auditory cognitive processing. This reorganization was linked to a general reduction in activity within memory-related brain regions, corroborating previous studies that have documented attenuated brain responses in older adults across various conditions, including resting-state activity, automatic neural responses, and consciously performed tasks [46]. Building upon these findings, the current study further extends the characterization of age-related differences by incorporating resting-state data into the analysis, thereby demonstrating age-related brain network reorganization both during task engagement and at rest. Moreover, the methodology adopted in the present study enabled a frequency-resolved analysis, yielding insights into age-related network differences across distinct frequency bands. The methodology adopted in the present study corresponds to the FREQ-NESS pipeline introduced in [47]. In such pipeline, GED was applied to source-reconstructed MEG recordings acquired during passive listening and resting state. This approach demonstrated efficacy in analyzing datasets involving continuous stimuli, such as music or speech, to reveal neural responses and connectivity patterns [47]. Here, the same FREQ-NESS analytical framework was extended to MEG data collected during an active memory task, thereby demonstrating the applicability of the method to extract relevant information from a more engaging cognitive paradigm compared to passive listening. Specifically, this approach enabled a time-resolved investigation of frequency dynamics; the application of the Morlet wavelet transform facilitated the identification of not only the dominant frequencies implicated in the task-related processes but also the temporal evolution of their significance and prominence.

The ANOVA test performed on the first component of variance explained at each frequency revealed significant effects for both condition and the interaction between age and condition at 1.07 Hz. This frequency falls within the delta range, which prior literature has associated with motor delta oscillations playing a key functional role in human auditory perception, both enabling and constraining the temporal flow of information [73]. The presence of delta-band activity during auditory tasks is well-established in the literature. Prior research suggests that motor delta oscillations play a critical role in auditory perception by both facilitating and constraining the temporal organization of sensory input. These oscillations contribute to the encoding of temporal context, thereby modulating auditory processing and influencing behavioral performance [73]. In particular, studies investigating melody recognition tasks have identified 1 Hz and 4 Hz as primary contributors to the MEG signal during such tasks [74]. Although a higher peak was observed in the listening condition for the younger adult group, the statistical analysis did not identify age as a significant factor; this aligns with existing research, which has reported few significant correlations between age and spectral characteristics of delta rhythms [75]. One study, which included patients with MCI and healthy controls, found that healthy controls exhibited a larger delta response compared to MCI patients during an auditory oddball paradigm [76]. While the present study did not address pathological conditions but focused instead on healthy aging, the observed reduction in the 1.07 Hz peak in older adults compared to younger ones appears promising for investigations into both healthy and pathological aging.

The significance identified by the ANOVA test at the stimulation frequency, 2.86 Hz, aligns with findings reported in [47]. In that study, a prominent peak at the stimulation frequency was observed in the variance explained, indicating a strong attunement of the brain to the frequency of auditory stimulation. “Attunement” here refers to the phenomenon where the spectral content of a network component becomes highly concentrated around a specific frequency as a direct effect of the stimulation [47]. While the stimulation context in that study involved passive listening, the present findings demonstrate a similar effect during an active melody-recognition task.

Another frequency found to be significant is 10.00 Hz, which lies within the alpha range. Notably, it is the interaction between age and condition that reached statistical significance. This observation aligns with existing literature reporting age-related changes in alpha-band activity during rest. Specifically, numerous studies have documented a decline in alpha-band parameters with aging [77]. A well-documented characteristic of EEG aging is the slowing of alpha rhythms with increasing age [75, 78, 79]. Comparative studies of alpha peak frequency between younger and older groups have shown that the latter group exhibits significantly lower frequencies [75, 78]. This pattern is consistent with the present study’s variance explained as a function of frequency: during resting state, the alpha-range peak appears at a lower frequency for older compared to younger adults. Correspondingly, several lifespan studies have reported that alpha peak frequency, especially in the posterior regions, increases linearly up to around 20 years of age and begins to decline around the age of 40–50 [80, 81]. Interestingly, the alpha peak in older adults increased during the task compared to younger participants. This suggests that age-related neurophysiological changes are not merely characterized by a reduced prominence of the alpha network, but rather by its remodulation depending on the context, such as engaging in a task versus simply resting

A further critical aging-related characteristic of the alpha rhythm is alpha reactivity. This refers, particularly in posterior regions, to a substantial reduction in alpha activity in response to visual stimuli or during cognitively demanding tasks that require focused attention or mental effort [75]. Alpha suppression is interpreted as a marker of active sensory information processing, promoting task-relevant activity and enabling other neural frequencies to dominate. Previous studies have documented reduced alpha reactivity in older compared to younger adults [82]. Although this study did not explicitly compute alpha reactivity, the observed results are consistent with this literature: while younger individuals show higher variance explained in the alpha band during resting state compared to task, this difference is less pronounced in older adults. Furthermore, for older adults, the variance explained during the task exceeds that observed during rest. These findings highlight the potential for future research to explore alpha reactivity more rigorously using the current approach, which permits detailed frequency-specific analysis. Such investigations could elucidate differences in alpha reactivity between young and older individuals, and potentially distinguish between healthy and pathological aging.

Brain networks time-frequency analysis revealed a significant difference between the resting state and task conditions within the frequency range of 7.10 to 22.90 Hz, corresponding to the alpha and beta bands, following stimulus onset. This result is consistent with prior research, which has associated post-stimulus desynchronization in these frequency bands with successful memory performance. In this context, event-related synchronization and desynchronization refer to power changes relative to a resting or prestimulus baseline, manifesting as either an increase (synchronization) or a decrease (desynchronization) in spectral power. Both alpha and beta bands typically show increased desynchronization approximately 600 ms post-stimulus onset, persisting throughout the epoch duration, in agreement with the present findings (stimulus onset occurred at 0.1 s) [83]. Additionally, beta power has been shown to exhibit strong associations with various brain networks, including positive correlations with the resting state [75], thereby supporting the observation of significant differences in the beta range between the listening and resting conditions.

Numerous studies have also explored the impact of aging on beta rhythms. Although some discrepancies exist in the literature [75], the prevailing view suggests an age-related increase in beta power; however, other findings [84] have reported a significant reduction in absolute beta power in the midline and occipital regions in older adults. In the present study, fewer and comparatively weaker effects (relative to the condition effect) were identified with respect to age and the interaction between age and condition, with most significant findings emerging in the beta frequency range.

The results presented here revealed significant differences between younger and older individuals. Conducted with a focus on healthy aging, the study successfully identified brain networks associated with specific frequencies that vary with age. A natural extension of this work would involve applying the same analytical framework to compare healthy and pathological older individuals, with the aim of distinguishing features specific to pathological aging. Notably, the standard error of the variance explained was generally higher among older participants, particularly in the higher frequency ranges (gamma), indicating greater inter-individual variability. This observation supports the idea of further stratifying the older cohort into narrower age groups to better characterize age-related neural changes over time. While previous studies have investigated age-related variations in specific neural features, they typically rely on conventional frequency bands. In contrast, the approach employed here enables a fine-grained, frequency-resolved investigation of brain networks and their evolution with age.

This enhanced spectral resolution could also be leveraged to examine cross-frequency coupling in greater detail. For instance, attentional modulation of auditory processing has been linked to top-down temporal predictions, which are at least partially generated in the motor cortex. These predictions are represented through delta–beta phase-amplitude coupling, where both delta phase and beta amplitude have been shown to modulate auditory responses and predict behavioral outcomes [73]. The present method offers the potential to investigate this coupling with greater spectral precision, by analyzing narrower frequency ranges. Moreover, given the relevance of delta–beta coupling in auditory perception, future studies could employ this approach to explore how such interactions vary with age and between healthy and pathological populations.

Finally, the observed age-related differences in alpha activity, where younger individuals exhibit higher variance explained during resting state and older individuals during the auditory memory task, suggest the potential for targeted investigations of alpha reactivity. The frequency resolution enabled by the current method would be particularly advantageous for such studies.

The consistency of the present findings with existing literature supports the validity of applying the FREQ-NESS pipeline [47], for the identification of functional brain networks during an active memory task. The analysis of variance explained as a function of frequency provided insight into the relevance of each frequency band, highlighting its specific contribution, and the statistical significance of that contribution, to auditory and memory processing. The examination of induced responses leveraged the high temporal resolution of MEG, offering valuable information on the temporal evolution of frequency-specific activity and the dynamic involvement of each frequency band in stimulus processing. In addition, only a limited number of significant differences were observed between networks with respect to condition or age. Indeed, only three frequency bands reached statistical significance. This result highlights the strength of the approach, as it indicates that a limited number of frequencies were of key importance in this context, thereby emphasizing the relevance of conducting frequency-specific brain network analysis. To enhance frequency resolution, future work could further refine the filtering procedure, thereby optimizing the extraction of frequency-resolved information. Aside from this common consideration regarding filter design, the FREQ-NESS pipeline applied to event-related source reconstructed MEG data demonstrated robustness and utility in capturing functionally meaningful brain dynamics during an active memory task.

## Code availability

The code for the preprocessing is available at the following link: https://github.com/leonardob92/LBPD-1.0.git

The pipeline for the current study is available at the following link: https://github.com/malvasochiara/FREQ-NESS-for-age-related-differences-in-brain-networks

The FREQ-NESS Toolbox is available at the following link: https://github.com/mattiaRosso92/Frequency-resolved_brain_network_estimation_via_source_separation_FREQ-NESS/tree/main/FREQNESS_Toolbox

## Acknowledgements

The Center for Music in the Brain (MIB) is funded by the Danish National Research Foundation (project number DNRF117). Additionally, we thank the Fundación Mutua Madrileña for the economic support provided to author GFR.

LB is supported by Independent Research Fund Denmark (DFF) Sapere Aude, Lundbeck Foundation (Talent Prize 2022), Carlsberg Foundation (CF20-0239), Center for Music in the Brain, Linacre College of the University of Oxford, Society for Education and Music Psychology (SEMPRE’s 50th Anniversary Awards Scheme), and Nordic Mensa Fund.

MLK is supported by Center for Music in the Brain and Centre for Eudaimonia and Human Flourishing, which is funded by the Pettit and Carlsberg Foundations.

## Author contributions

LB, CM, MR and GFR conceived the hypotheses. LB and GFR designed the study. LB, MLK, CT, ES, VR and PV recruited the resources for the experiment. LB and GFR collected the data. CM, MR and LB performed pre-processing, statistical analysis and developed the adapted FREQ-NESS pipeline. MLK, PV, ES, CT and VR provided essential help to interpret and frame the results within the neuroscientific and methodological literature. CM, with the contribution from LB, wrote the first draft of the manuscript. GFR, CM and LB prepared the figures. All the authors contributed to and approved the final version of the manuscript.

## References

[1] Ian J. Deary et al. “Age-associated cognitive decline”. In: British Medical Bulletin 92.1 (Sept. 2009), pp. 135–152. issn: 0007-1420. doi: 10.1093/bmb/ldp033. eprint: https://academic.oup.com/bmb/article-pdf/92/1/135/951616/ldp033.pdf. url: https://doi.org/10.1093/bmb/ldp033.

[2] Raymond Levy. “Aging-Associated Cognitive Decline”. In: International Psychogeriatrics 6.1 (1994), pp. 63–68. doi: 10.1017/S1041610294001626.

[3] Gemma Fernández-Rubio et al. “Investigating the impact of age on auditory short-term, long-term, and working memory”. In: Psychology of Music 52.2 (2024), pp. 187–198. doi: 10.1177/03057356231183404. eprint: https://doi.org/10.1177/03057356231183404. url: https://doi.org/10.1177/03057356231183404.

[4] L. Bonetti et al. “Working Memory Predicts Long-Term Recognition of Auditory Sequences: Dissociation Between Confirmed Predictions and Prediction Errors”. In: Scandinavian Journal of Psychology (May 2025). Open Access; First published: 21 May 2025. doi: 10.1111/sjop.13124. url: https://doi.org/10.1111/sjop.13124.

[5] Macarena Sánchez-Izquierdo and Rocío Fernández-Ballesteros. “Cognition in Healthy Aging”. In: International Journal of Environmental Research and Public Health 18.3 (2021). issn: 1660-4601. doi: 10.3390/ijerph18030962. url: https://www.mdpi.com/1660-4601/18/3/962.

6. Gemma Fernández-Rubio, et al. “The neurophysiology of healthy and pathological aging: A comprehensive systematic review”. In: bioRxiv (2024). doi: 10.1101/2024.08.06.606817. eprint: https://www.biorxiv.org/content/early/2024/08/07/2024.08.06.606817. full.pdf. url: https://www.biorxiv.org/content/early/2024/08/07/2024.08.06.606817.

[7] C. Babiloni et al. “Brain neural synchronization and functional coupling in Alzheimer’s disease as revealed by resting state EEG rhythms”. In: International Journal of Psychophysiology 103 (2016), pp. 88–102. doi: 10.1016/j.ijpsycho.2015.02.008.

[8] Y. Ding, Y. Chu, M. Liu, et al. “Fully automated discrimination of Alzheimer’s disease using resting-state electroencephalography signals”. In: Quantitative Imaging in Medicine and Surgery 12.2 (2022), pp. 1063–1078. doi: 10.21037/qims-21-430.

[9] Saúl J. Ruiz-Gómez et al. “Automated Multiclass Classification of Spontaneous EEG Activity in Alzheimer’s Disease and Mild Cognitive Impairment”. In: Entropy 20.1 (2018). issn: 1099-4300. doi: 10.3390/e20010035. url: https://www.mdpi.com/1099-4300/20/1/35.

[10] C. Babiloni et al. “Resting State Alpha Electroencephalographic Rhythms Are Differently Related to Aging in Cognitively Unimpaired Seniors and Patients with Alzheimer’s Disease and Amnesic Mild Cognitive Impairment”. In: Journal of Alzheimer’s Disease 82.3 (2021), pp. 1085–1114. doi: 10.3233/JAD-201271.

[11] G. Lu et al. “Neuroimaging of EEG Rhythms at Resting State in Normal Elderly Adults: A Standard Low-Resolution Electromagnetic Tomography Study”. In: Journal of Clinical Neurophysiology 39.1 (2022), pp. 72–77. doi: 10.1097/WNP.0000000000000780.

[12] B. Scally et al. “Resting-state EEG power and connectivity are associated with alpha peak frequency slowing in healthy aging”. In: Neurobiology of Aging 71 (2018), pp. 149–155. doi: 10.1016/j.neurobiolaging.2018.07.004.

[13] Fernando Maestú and Alberto Fernández. “Role of Magnetoencephalography in the Early Stages of Alzheimer Disease”. In: Neuroimaging Clinics of North America 30.2 (2020). Magnetoencephalography, pp. 217–227. issn: 1052-5149. doi: 10.1016/j.nic.2020.01.003. url: https://www.sciencedirect.com/science/article/pii/S1052514920300034.

[14] J. Dauwels, F. Vialatte, and A. Cichocki. “Diagnosis of Alzheimer’s disease from EEG signals: where are we standing?” In: Current Alzheimer Research 7.6 (2010), pp. 487–505. doi: 10.2174/156720510792231720.

[15] D. Puttaert, V. Wens, P. Fery, et al. “Decreased Alpha Peak Frequency Is Linked to Episodic Memory Impairment in Pathological Aging”. In: Frontiers in Aging Neuroscience 13 (Aug. 2021). Published 2021 Aug 12, p. 711375. doi: 10.3389/fnagi.2021.711375.

[16] D. López-Sanz et al. “Alpha band disruption in the AD-continuum starts in the Subjective Cognitive Decline stage: a MEG study”. In: Scientific Reports 6 (2016), p. 37685. doi: 10.1038/srep37685.

[17] R. Cassani et al. “Systematic Review on Resting-State EEG for Alzheimer’s Disease Diagnosis and Progression Assessment”. In: Disease Markers 2018 (2018). Published 2018 Oct 4, p. 5174815. doi: 10.1155/2018/5174815.

[18] R. J. Barry et al. “EEG differences between eyes-closed and eyesopen resting conditions”. In: Clinical Neurophysiology 118.12 (2007), pp. 2765–2773. doi: 10.1016/j.clinph.2007.07.028.

[19] Pin-Yu Chen et al. “Altered mismatch response of inferior parietal lobule in amnestic mild cognitive impairment: A magnetoencephalographic study”. In: CNS Neuroscience & Therapeutics 27.10 (2021), pp. 1136–1145. doi: 10.1111/cns.13691. url: https://onlinelibrary.wiley.com/doi/epdf/10.1111/cns.13691.

[20] Loren Mowszowski et al. “Reduced mismatch negativity in mild cognitive impairment: associations with neuropsychological performance”. In: Journal of Alzheimer’s Disease 30.1 (2012), pp. 209–219. doi: 10.3233/JAD-2012-111868. url: https://content.iospress.com/doi/10.3233/JAD-2012-111868.

[21] Shixiang Jiang et al. “Mismatch negativity as a potential neurobiological marker of early-stage Alzheimer disease and vascular dementia”. In: Neuroscience Letters 647 (2017), pp. 26–31. doi: 10.1016/j.neulet.2017.03.032. url: https://www.sciencedirect.com/science/article/pii/S0304394017302525.

[22] Yana Criel et al. “Aging and sex effects on phoneme perception: An exploratory mismatch negativity and P300 investigation”. In: International Journal of Psychophysiology 190 (2023), pp. 69–83. doi: 10.1016/j.ijpsycho.2023.06.002. url: 10.1016/j.ijpsycho.2023.06.002.

[23] Michael A. Kisley et al. “Age-related change in neural processing of time-dependent stimulus features”. In: Cognitive Brain Research 25.3 (2005), pp. 913–925. doi: 10.1016/j.cogbrainres.2005.09.014. url: 10.1016/j.cogbrainres.2005.09.014.

[24] Elissa-Marie Cocquyt et al. “Effects of Healthy Aging and Gender on the Electrophysiological Correlates of Semantic Sentence Comprehension: The Development of Dutch Normative Data”. In: Journal of Speech, Language, and Hearing Research 66.5 (2023), pp. 1694–1717. doi: 10.1044/2023_JSLHR-22-00545. url:10.1044/2023_JSLHR-22-00545.

[25] Hannes O. Tiedt, Felicitas Ehlen, and Fabian Klostermann. “Age-related dissociation of N400 effect and lexical priming”. In: Scientific Reports 10.1 (2020), p. 20200. doi: 10.1038/s41598-020-77116-9. url: https://www.nature.com/articles/s41598-020-77116-9.

[26] Nannan Xu et al. “Increased world knowledge in older adults does not prevent decline in world knowledge comprehension: An ERP study”. In: Brain and Cognition 140 (2020), p. 105534. doi: 10.1016/j.bandc.2020.105534. url: 10.1016/j.bandc.2020.105534.

[27] Marilyne Joyal et al. “Semantic Processing in Healthy Aging and Alzheimer’s Disease: A Systematic Review of the N400 Differences”. In: Brain Sciences 10.11 (2020), p. 770. doi: 10.3390/brainsci10110770. url: https://www.mdpi.com/2076-3425/10/11/770.

[28] Kathleen H. Elverman et al. “Event-Related Potentials, Inhibition, and Risk for Alzheimer’s Disease Among Cognitively Intact Elders”. In: Journal of Alzheimer’s Disease 80.4 (2021), pp. 1413–1428. doi: 10.3233/JAD-201559. url: https://content.iospress.com/doi/10.3233/JAD-201559.

[29] Gemma Learmonth et al. “Age-related reduction of hemispheric lateralisation for spatial attention: An EEG study”. In: NeuroImage 153 (2017), pp. 139–151. doi: 10.1016/j.neuroimage.2017.03.050. url: 10.1016/j.neuroimage.2017.03.050.

[30] H. Burianová et al. “Motor neuroplasticity: A MEG-fMRI study of motor imagery and execution in healthy ageing”. In: Neuropsychologia 146 (2020), p. 107539. doi: 10.1016/j.neuropsychologia.2020.107539.

[31] G. Zhang et al. “Computational exploration of dynamic mechanisms of steady state visual evoked potentials at the whole brain level”. In: Neuroimage 237 (2021), p. 118166. doi: 10.1016/j.neuroimage.2021.118166.

[32] H. E. Rossiter et al. “Beta oscillations reflect changes in motor cortex inhibition in healthy ageing”. In: Neuroimage 91 (2014), pp. 360–365. doi: 10.1016/j.neuroimage.2014.01.012.

[33] Antonio Criscuolo et al. “Aging Impacts Basic Auditory and Timing Processes”. In: European Journal of Neuroscience (Mar. 2025). Open Access; Associate Editor: Yoland Smith. doi: 10.1111/ejn.70031. url: 10.1111/ejn.70031.

[34] P. Vuust et al. “Music in the brain”. In: Nature Reviews Neuroscience 23.5 (2022), pp. 287–305. doi: 10.1038/s41583-022-00578-5.

[35] L. Bonetti et al. “Rapid encoding of musical tones discovered in whole-brain connectivity”. In: Neuroimage 245 (2021), p. 118735. doi: 10.1016/j.neuroimage.2021.118735.

[36] L. Bonetti et al. “Spatiotemporal brain hierarchies of auditory memory recognition and predictive coding”. In: Nature Communications 15.1 (2024). Published 2024 May 21, p. 4313. doi: 10.1038/s41467-024-48302-4.

[37] M. Costa et al. “EEG Correlates of Auditory Short-Term Memory and Dissimilarity Perception in Young and Older Adults”. In: European Journal of Neuroscience (June 2025). Open Access; First published: 17 June 2025; Associate Editor: Edmund Lalor. doi: 10.1111/ejn.70166. url: 10.1111/ejn.70166.

[38] Ramón Nartallo-Kaluarachchi et al. “Multilevel Irreversibility Reveals Higher-Order Organization of Nonequilibrium Interactions in Human Brain Dynamics”. In: Proceedings of the National Academy of Sciences 122.10 (Mar. 2025). Edited by Danielle Basset; accepted by Elizabeth A. Buffalo on January 28, 2025, e2408791122. doi: 10.1073/pnas.2408791122. url: 10.1073/pnas.2408791122.

[39] L. Bonetti, et al. “The neural mechanisms of concept formation over time in music”. In: bioRxiv (2024). doi: 10.1101/2024.11.06.622228. eprint: https://www.biorxiv.org/content/early/2024/11/07/2024.11.06.622228.full.pdf. url: https://www.biorxiv.org/content/early/2024/11/07/2024.11.06.622228.

[40] David R. Quiroga-Martinez et al. “Decoding Reveals the Neural Representation of Perceived and Imagined Musical Sounds”. In: PLOS Biology (Oct. 2024). Version 2, Published: October 21, 2024, e3002858. doi: 10.1371/journal.pbio.3002858. url: 10.1371/journal.pbio.3002858.

[41] Leonardo Bonetti et al. “Spatiotemporal whole-brain activity and functional connectivity of melodies recognition”. In: Cerebral Cortex 34.8 (Aug. 2024), bhae320. issn: 1460-2199. doi: 10.1093/cercor/bhae320. eprint: https://academic.oup.com/cercor/article-pdf/34/8/bhae320/58756359/bhae320.pdf. url: 10.1093/cercor/bhae320.

[42] Steffen A. Herff et al. “Hierarchical syntax model of music predicts theta power during music listening”. In: Neuropsychologia 199 (2024), p. 108905. issn: 0028-3932. doi: 10.1016/j.neuropsychologia.2024.108905. url: https://www.sciencedirect.com/science/article/pii/S0028393224001209.

[43] Leonardo Bonetti, et al. “BROAD-NESS uncovers dual-stream mechanisms underlying predictive coding in auditory memory networks”. In: bioRxiv (2025). doi: 10.1101/2024.10.31.621257. eprint: https://www.biorxiv.org/content/early/2025/03/11/2024.10.31.621257.full.pdf. url: https://www.biorxiv.org/content/early/2025/03/11/2024.10.31.621257.

[44] E. Serra, et al. “Neurophysiological correlates of short-term recognition of sounds: Insights from magnetoencephalography”. In: bioRxiv (2023). doi: 10.1101/2023.12.07.570594. eprint: https://www.biorxiv.org/content/early/2023/12/08/2023.12.07.570594.full.pdf. url: https://www.biorxiv.org/content/early/2023/12/08/2023.12.07.570594.

[45] L Bonetti et al. “Brain recognition of previously learned versus novel temporal sequences: a differential simultaneous processing”. In: Cerebral Cortex 33.9 (Nov. 2022), pp. 5524–5537. issn: 1047-3211. doi: 10.1093/cercor/bhac439. eprint: https://academic.oup.com/cercor/article-pdf/33/9/5524/50096454/bhac439.pdf. url: 10.1093/cercor/bhac439.

[46] L. Bonetti, G. Fernández-Rubio, and M. Lumaca. “Age-related neural changes underlying long-term recognition of musical sequences”. In: Communications Biology 7 (2024), p. 1036. doi: 10.1038/s42003-024-06587-7. url: 10.1038/s42003-024-06587-7.

[47] M. Rosso, G. Fernández-Rubio, P.E. Keller, et al. “FREQ-NESS Reveals the Dynamic Reconfiguration of Frequency-Resolved Brain Networks During Auditory Stimulation”. In: Advanced Science (Weinheim) (2025). Published online April 10, 2025. doi: 10.1002/advs.202413195.

[48] Michael X Cohen. “A tutorial on generalized eigendecomposition for denoising, contrast enhancement, and dimension reduction in multichannel electrophysiology”. In: NeuroImage 247 (2022), p. 118809. issn: 1053-8119. doi: 10.1016/j.neuroimage.2021.118809. url: https://www.sciencedirect.com/science/article/pii/S1053811921010806.

[49] Mattia Rosso et al. “Neural entrainment underpins sensorimotor synchronization to dynamic rhythmic stimuli”. In: NeuroImage 277 (June 2023), p. 120226. doi: 10.1016/j.neuroimage.2023.120226.

[50] Mattia Rosso et al. “Mutual beta power modulation in dyadic entrainment”. In: NeuroImage 257 (Aug. 2022), p. 119326. doi: 10.1016/j.neuroimage.2022.119326.

[51] Mattia Rosso, Marc Leman, and Lousin Moumdjian. “Neural Entrainment Meets Behavior: The Stability Index as a Neural Outcome Measure of Auditory-Motor Coupling”. In: Frontiers in Human Neuroscience Volume 15 - 2021 (2021). issn: 1662-5161. doi: 10.3389/fnhum.2021.668918. url: https://www.frontiersin.org/journals/human-neuroscience/articles/10.3389/fnhum.2021.668918.

[52] M. B. Zuure et al. “Multiple Midfrontal Thetas Revealed by Source Separation of Simultaneous MEG and EEG”. In: Journal of Neuroscience 40.40 (2020), pp. 7702–7713. doi: 10.1523/JNEUROSCI.0321-20.2020.

[53] Alain Cheveigné and Dorothée Arzounian. “Scanning for oscillations”. In: Journal of Neural Engineering 12 (Oct. 2015), p. 066020. doi: 10.1088/1741-2560/12/6/066020.

[54] Stefan Haufe et al. “On the interpretation of weight vectors of linear models in multivariate neuroimaging”. In: NeuroImage 87 (2014), pp. 96–110. issn: 1053-8119. doi: 10.1016/j.neuroimage.2013.10.067. url: https://www.sciencedirect.com/science/article/pii/S1053811913010914.

[55] L Bonetti et al. “Brain recognition of previously learned versus novel temporal sequences: a differential simultaneous processing”. In: Cerebral Cortex 33.9 (Nov. 2022), pp. 5524–5537. issn: 1047-3211. doi: 10.1093/cercor/bhac439. eprint: https://academic.oup.com/cercor/article-pdf/33/9/5524/50096454/bhac439.pdf. url: 10.1093/cercor/bhac439.

[56] G. Fernández-Rubio, E. Brattico, S.A. Kotz, et al. “Magnetoencephalography recordings reveal the spatiotemporal dynamics of recognition memory for complex versus simple auditory sequences”. In: Communications Biology 5 (2022), p. 1272. doi: 10.1038/s42003-022-04217-8.

[57] G. Fernández-Rubio et al. “Associations between abstract working memory abilities and brain activity underlying long-term recognition of auditory sequences”. In: PNAS Nexus 1.4 (2022), pgac216. doi: 10.1093/pnasnexus/pgac216.

58. [58] L. Bonetti, et al. “Spatiotemporal brain dynamics during recognition of the music of Johann Sebastian Bach”. In: bioRxiv (2020). doi: 10.1101/2020.06.23.165191. eprint: https://www.biorxiv.org/content/early/2020/06/24/2020.06.23.165191.full.pdf. url: https://www.biorxiv.org/content/early/2020/06/24/2020.06.23.165191.

[59] L. Bonetti, G. Fernández-Rubio, F. Carlomagno, et al. “Spatiotemporal brain hierarchies of auditory memory recognition and predictive coding”. In: Nature Communications 15 (2024), p. 4313. doi: 10.1038/s41467-024-48302-4.

[60] S. Taulu and J. Simola. “Spatiotemporal signal space separation method for rejecting nearby interference in MEG measurements”. In: Physics in Medicine and Biology 51.7 (2006), pp. 1759–1768. doi: 10.1088/0031-9155/51/7/008.

[61] M. Woolrich et al. “MEG beamforming using Bayesian PCA for adaptive data covariance matrix regularization”. In: Neuroimage 57.4 (2011), pp. 1466–1479. doi: 10.1016/j.neuroimage.2011.04.041.

[62] Oostenveld R. et al. “FieldTrip: Open source software for advanced analysis of MEG, EEG, and invasive electrophysiological data”. In: Comput Intell Neurosci 2011 (2011), p. 156869. doi: 10.1155/2011/156869.

[63] Mark W. Woolrich et al. “Bayesian analysis of neuroimaging data in FSL”. In: NeuroImage 45.1, Supplement 1 (2009). Mathematics in Brain Imaging, S173–S186. issn: 1053-8119. doi: 10.1016/j.neuroimage.2008.10.055. url: https://www.sciencedirect.com/science/article/pii/S1053811908012044.

[64] W. D. Penny et al. Statistical parametric mapping: the analysis of functional brain images. Elsevier, 2011.

[65] Mantini D. et al. “A signal-processing pipeline for magnetoencephalography resting-state networks”. In: Brain Connect 1.1 (2011), pp. 49–59. doi: 10.1089/brain.2011.0001.

[66] G. Nolte. “The magnetic lead field theorem in the quasi-static approximation and its use for magnetoencephalography forward calculation in realistic volume conductors”. In: Physics in Medicine and Biology 48.22 (2003), pp. 3637–3652. doi: 10.1088/0031-9155/48/22/002.

[67] M.X. Huang, J.J. Shih, R.R. Lee, et al. “Commonalities and differences among vectorized beamformers in electromagnetic source imaging”. In: Brain Topography 16.3 (2004), pp. 139–158. doi: 10.1023/b:brat.0000019183.92439.51.

[68] M.X. Huang, J.C. Mosher, and R.M. Leahy. “A sensor-weighted overlapping-sphere head model and exhaustive head model comparison for MEG”. In: Physics in Medicine and Biology 44.2 (1999), pp. 423–440. doi: 10.1088/0031-9155/44/2/010.

[69] Alex Pappachen James, Aidyn Zhambyl, and Anju Nandakumar. Memristor-based Approximation of Gaussian Filter. 2018. arXiv: 1805 . 06643 [eess.SP]. url: https://arxiv.org/abs/1805.06643.

[70] L. Bonetti, et al. “BROADband brain Network Estimation via Source Separation (BROAD-NESS)”. In: bioRxiv (2024). doi: 10.1101/2024.10.31.621257. eprint: https://www.biorxiv.org/content/early/2024/10/31/2024.10.31.621257.full.pdf. url: https://www.biorxiv.org/content/early/2024/10/31/2024.10.31.621257.

[71] Michael X Cohen. “A better way to define and describe Morlet wavelets for time-frequency analysis”. In: NeuroImage 199 (2019), pp. 81–86. issn: 1053-8119. doi: 10.1016/j.neuroimage.2019.05.048. url: https://www.sciencedirect.com/science/article/pii/S1053811919304409.

[72] Andreas Bruns. “Fourier-, Hilbert- and wavelet-based signal analysis: are they really different approaches?” In: Journal of Neuroscience Methods 137.2 (2004), pp. 321–332.

[73] Benjamin Morillon et al. “Prominence of delta oscillatory rhythms in the motor cortex and their relevance for auditory and speech perception”. In: Neuroscience & Biobehavioral Reviews 107 (2019), pp. 136–142. issn: 0149-7634. doi: 10.1016/j.neubiorev.2019.09.012. url: https://www.sciencedirect.com/science/article/pii/S0149763419300922.

[74] Leonardo Bonetti et al. “Spatiotemporal whole-brain activity and functional connectivity of melodies recognition”. In: Cerebral Cortex 34.8 (Aug. 2024), bhae320.

[75] Jae-Hwan Kang, Jang-Han Bae, and Young-Ju Jeon. “Age-Related Characteristics of Resting-State Electroencephalographic Signals and the Corresponding Analytic Approaches: A Review”. In: Bioengineering 11.5 (Apr. 2024). Ed. by Yunfeng Wu, p. 418. doi: 10.3390/bioengineering11050418.

[76] Pınar Kurt et al. “Patients with Mild Cognitive Impairment Display Reduced Auditory Event-Related Delta Oscillatory Responses”. In: Brain Research 1587 (2014), pp. 79–89. doi: 10.1016/j.brainres.2014.04.025.

[77] Cécile Bordier et al. “Age-Related Differences in Resting-State EEG and Allocentric Spatial Working Memory Performance”. In: Frontiers in Aging Neuroscience 13 (Nov. 2021). doi: 10.3389/fnagi.2021.704362. url: https://www.frontiersin.org/articles/10.3389/fnag.

[78] Jemaine E. Stacey et al. “Age differences in resting state EEG and their relation to eye movements and cognitive performance”. In: Neuropsychologia 157 (July 2021), p. 107887. doi: 10.1016/j.neuropsychologia.2021.107887.

[79] Brian Scally et al. “Resting-state EEG power and connectivity are associated with alpha peak frequency slowing in healthy aging”. In: Neurobiology of Aging 71 (Nov. 2018), pp. 149–155. doi: 10.1016/j.neurobiolaging.2018.07.004.

[80] A. K. I. Chiang et al. “Age trends and sex differences of alpha rhythms including split alpha peaks”. In: Clinical Neurophysiology 122.8 (Aug. 2011), pp. 1505–1517. doi: 10.1016/j.clinph.2011.01.040.

[81] H. Aurlien et al. “EEG background activity described by a large computerized database”. In: Clinical Neurophysiology 115.3 (Mar. 2004), pp. 665–673. doi: 10.1016/j.clinph.2003.10.019.

[82] N. V. Volf and A. A. Gluhih. “Background Cerebral Electrical Activity in Healthy Mental Aging”. In: Human Physiology 37.4 (2011). Research Institute of Physiology, Siberian Branch, Russian Academy of Medical Sciences, Novosibirsk, Russia, pp. 418–423. doi: 10.1134/S0362119711040207.

[83] J. Strunk et al. “Age-related changes in neural oscillations supporting context memory retrieval”. In: Cortex 91 (2017), pp. 40–55. doi: 10.1016/j.cortex.2017.01.020.

[84] J. Breslau et al. “Topographic EEG changes with normal aging and SDAT”. In: Electroencephalography and Clinical Neurophysiology 72.4 (1989), pp. 281–289. doi: 10.1016/0013-4694(89)90063-1.

